# Dynamic Functional Network Connectivity In Schizophrenia With MEG And fMRI, Do Different Time Scales Tell A Different Story?

**DOI:** 10.1101/432385

**Authors:** Lori Sanfratello, Jon Houck, Vince Calhoun

## Abstract

The importance of how brain networks function together to create brain states has become increasingly recognized. Therefore, an investigation of eyes-open resting state dynamic functional network connectivity (dFNC) of healthy controls (HC) versus that of schizophrenia patients (SP) via both fMRI and a novel MEG pipeline was completed. The fMRI analysis used a spatial independent component analysis (ICA) to determine the networks on which the dFNC was based. The MEG analysis utilized a source-space activity estimate (MNE/dSPM) whose result was the input to a spatial ICA, on which the networks of the MEG dFNC was based. We found that dFNC measures reveal significant differences between HC and SP, which depended upon the imaging modality. Consistent with previous findings, a dFNC analysis predicated on fMRI data revealed HC and SP remain in different overall brain states (defined by a k-means clustering of network correlations) for significantly different periods of time, with SP spending less time in a highly-connected state. The MEG dFNC, in contrast, revealed group differences in more global statistics: SP changed between meta-states (k-means cluster states that are allowed to overlap in time) significantly more often and to states which were more different, relative to HC. MEG dFNC also revealed a highly connected state where a significant difference was observed in inter-individual variability, with greater variability among SP. Overall, our results show that fMRI and MEG reveal between-group functional connectivity differences in distinct ways, highlighting the utility of using each of the modalities individually, or potentially a combination of modalities, to better inform our understanding of disorders such as schizophrenia.

## 1. Introduction

Noninvasive neuroimaging is currently available in a number of modalities, including functional MRI (fMRI) with its excellent spatial resolution, and magnetoencephalography (MEG) with its excellent temporal resolution. Harnessing and/or combining the strengths these modalities offer is of great interest in both healthy and patient populations, which can aid in informing the classification of patient populations, identification of new treatment targets, and/or potential treatment success. One of the main goals of multi-modal imaging is to provide clinicians with biomarkers to assist with producing a diagnosis with increased confidence, and predicting a long-term prognosis for each new patient. Importantly, theoretical and experimental evidence imply that the biological signals detected by both fMRI and MEG originate from post-synaptic currents, albeit in a complex manner potentially variable by brain region (Ahonen et al., 1993; Hamalainen et al., 1993; Conner et al., 2011; Zhu et al., 2009; Harvey et al., 2013), indicating that the combination and/or comparison of MEG and fMRI makes theoretical sense (Hall et al., 2014). Although the exact relationship and influence of particular frequency bands remains an open question, the relationship between BOLD signal and electrophysiological activation has been shown for a variety of task activations in gamma band (Niessing, et al., 2005; Lachaux et al., 2007; Zaehle et al., 2009; Scheeringa et al., 2011; Kunii et al,. 2013), and it has been shown that in the resting state the direction (positive or negative) of EEG-BOLD signal correlations vary across brain regions and frequency bands, with lower as well as higher frequency brain oscillations linked to neurovascular processes. Of particular relevance here, it was found that low-frequency oscillations (<20 Hz), and not gamma activity, predominantly contributed to interareal BOLD correlations (Wang et al., 2012). The authors report that these low-frequency oscillations also influenced local processing by modulating gamma activity within individual areas, and suggest that such cross-frequency coupling links local BOLD signals to correlations across distributed networks. In addition, results from Bridwell et al. (2013) characterized brain networks spatially and spectrally, revealing that positive and negative associations appear within overlapping regions of the EEG frequency spectrum. That is, positive associations were primarily present within the lower (delta and theta) and higher (upper beta and lower gamma) spectral regions, sometimes within the same brain regions as measured by fMRI. Finally, it has been shown that even though the two modalities (fMRI and M/EEG) may exhibit activity in similar spatial locations, the functional pattern of this activity may differ in a complex manner, suggesting that each modality may be tuned to different aspects of neuronal activity (Muthukumaraswamy et al., 2008). Taken together these findings imply a complex relationship between neuronal activation and neurovascular coupling as measured by the BOLD signal, and suggest that all frequency bands contained in the M/EEG signal are potentially of interest when studying patient populations.

It has been shown in fMRI by Abrol et al. (2016) that multiple discrete, reoccurring connectivity states arise during rest, and that subjects tend to remain in one connectivity state for relatively long periods of time before transitioning to another. Other researchers observed how brain regions spontaneously changed their “module affiliations” (i.e., network connectivity) on a temporal scale of seconds, which could not be simply attributable to head motion or other error (Liao, et al., 2017, Vergara, et al., 2017). Similarly, in electrophysiological data it has been shown that sensor-space “microstates” arise and change on the order of hundreds of milliseconds (Van de Ville, et al., 2010; Khanna et al., 2015; Baker et al., 2014) and vary between disorders dependent upon which network a microstate correlated with (e.g. a microstate which correlated with the frontoparietal network was impaired in schizophrenia, Nishida et al., 2013). Patients with schizophrenia (SP) have been investigated in this manner by others as well, with SP experiencing particular microstates more often, and also experiencing briefer microstates than did healthy controls (HC)(review: Rieger et al., 2016).

One way to summarize the recurring connectivity states described above is via a functional network connectivity (FNC) analysis, which may be defined as the way in which sets of brain areas (networks) work together over time to produce different brain states, represented by statistical associations between network timecourses without regard to the spatial proximity of the regions to one another. Since an increasing body of literature suggests that neural oscillations perform a key role in binding separate brain regions together and promoting information transfer between distant brain areas (Buzsáki and Draguhn, 2004; Engel and Singer, 2001; Roopun et al., 2008) FNC has become an important metric for the study of how this naturally occurs (Jafri et al., 2008; Allen et al., 2014, Calhoun, et al., 2014). Furthermore, connectivity between brain regions is now generally accepted as being key to healthy brain function (Hall et al., 2014). Clearly the temporal as well as the spatial properties of these networks is of importance to our understanding of brain function, and it is probable that the definition of a network may vary on different time scales (Erhardt et al., 2011). Over the past decade FNC has most often been investigated within the resting state (i.e. in the absence of a defined task) using fMRI (Biswal, et al., 1995; Biswal et al., 2012, Allen et al., 2011) and, to a lesser extent electrophysiological methods, MEG and EEG (Brookes et al., 2011; Meier et al,. 2016; Nugent et al., 2016; Allen et al., 2017) in diverse populations including schizophrenia, depression, bipolar disorder, and in aging (Cetin et al., 2016; Houck et al., 2017; Wu et al., 2017; Fox, 2017; Du, et al., 2016; Nashiro et al., 2017; Madden 2017; Roiser, et al., 2016; Dong, et al., 2017; Alamian, et al., 2017). To date, FNC have most often been assessed as a static feature of the data, inferred from the overall (arbitrary) duration of the scan. But there is no a priori reason to assume that in the resting brain network correlations are static; indeed based on the known rapid dynamics of brain oscillations it may instead be expected that connections across networks change over time, even during a brief resting scan, as subjects experience different mental and emotional states (Miller et al., 2014 and 2016). The logical extension of FNC that looks at how states vary over small time “windows” in order to capture networks on a finer temporal scale, has been termed dynamic functional network connectivity (dFNC). dFNC has also been investigated in resting state fMRI (Sakoğlu et al., 2010; Miller, et al., 2014) and in diagnostic groups such as schizophrenia patients (Damaraju et al., 2014; Miller, et al., 2014) where for schizophrenia patients in an eyes-closed resting scan it has been shown that there is a reduction in fluidity, or dynamism, in their ability to move from state to state. Importantly, the dFNC spatial patterns of intra-subject dynamic variability have been shown to largely overlap with that of inter-subject variability, both of which were highly reproducible across repeated scanning sessions (Abrol et al., 2016). dFNC has therefore been established as a useful tool for both investigating changing brain states as well as for determining how these states vary between patient populations.

Here we present a dynamic functional network connectivity (dFNC) analysis using MEG and fMRI eyes-open resting scans collected from the same sample of healthy controls and schizophrenia patients, and indicate where we find overlap and differences between the results from the different modalities. We discuss possible reasons for these results, including careful selection of analysis parameters and scan length, particularly for the MEG data analysis where this has not been extensively studied previously. For the dFNC of fMRI data we follow an established pipeline (Calhoun et al., 2014; Miller et al., 2014). For the MEG data analysis, we describe the creation of a novel pipeline which includes a source-space analysis (MNE/dSPM, i.e. dynamic statistical parametric mapping from within Minimum Norm Estimate software) as input to a spatial ICA as the basis for the dFNC. We argue that using components calculated in such a way helps mitigate the “signal leakage” problem for MEG source-space based analyses as leakage manifests in the spatial maps, but the network connectivity patterns are preserved (Houck, et al., 2017). Finally, we investigate group differences between healthy controls and schizophrenia patients for both methods. We expect that SP will spend less time in highly connected cognitive states. We further hypothesize that for both fMRI and MEG analysis SP will show a reduction in global meta-state statistic values that measure how and when individuals move between states at a global level, relative to HC (i.e. we will see a “reduced dynamism” for SP).

## 2. Methods

### 2.1 Participants

Briefly, this investigation combined existing data (Aine et al., 2017) from 46 schizophrenia patients and 45 healthy controls from whom informed consent was obtained according to institutional guidelines at the University of New Mexico Human Research Protections Office (HRPO). All participants were compensated for their participation. Patients with a diagnosis of schizophrenia or schizoaffective disorder were invited to participate. Each patient completed the Structured Clinical Interview for DSM-IV Axis I Disorders (First et al., 1997) for diagnostic confirmation and evaluation of co-morbidities. Exclusion criteria included history of neurological disorders, mental retardation, substance abuse, or clinical instability. Patients were treated with a variety of antipsychotic medications, therefore doses of antipsychotic medications were converted to olanzapine equivalents (see Table 1: Gardner et al., 2010). Although patients and controls were not specifically matched, demographic characteristics including age, gender, and caregiver socio-economic status (Werner et al., 2007) were monitored throughout recruitment to ensure that both groups were of similar composition. There were no significant between-group differences on these measures. Each participant completed resting MEG and fMRI scans; however, only data from 74 participants was available to be used for the MEG analysis (demographics remained similar between groups, no significant differences were found). The participant data and preprocessing used for this study overlapped with that presented in Houck et al. (2017) but the analytic approach developed in this work is novel and distinct.

**Table 1.** ICA component numbers and anatomical locations

| **MEG ICA**<br>**Component #** | **Component ID** |
| --- | --- |
| 1 | Auditory (Left) |
| 2 | Posterior Cingulate (Bilat) |
| 3 | Lateral Occipital (Right) |
| 4 | Parahippocampus (Bilat) |
| 5 | Superior Frontal (Right) |
| 6 | Precuneus (Right) |
| 7 | Medial Orbital Frontal (Bilat) |
| 8 | Supramarginal (Left) |
| 9 | Temporal Pole (Right) |
| 10 | Insula (Right) |
| 11 | Visual (Left) |
| 12 | Precuneus (Left) |
| 13 | Paracentral (Right) |
| 14 | IFG/Insula/Lateral Orbital Front (Left) |
| 15 | Temporal Pole (Left) |
| 16 | Precuneus (Left) |
| 17 | Paracentral (Left) |
| 18 | Supramarginal (Right) |
| 19 | Frontal (Bilat) |
| 20 | Inferior Parietal (Right) |
| 21 | Postcentral (Right) |
| 22 | Insula (Left) |
| 23 | Inferior Parietal (Left) |
| 24 | Superior Frontal (Left) |
| 25 | Auditory (Right) |
| 26 | IFG/Insula/Lateral Orbital Front (Right) |
| 27 | Postcentral (Left) |
| 28 | Isthmus Cingulate (Bilat) |
| 29 | Lingual (Bilat) |
| 30 | MTG (Left) |
| 31 | MTG (Right) |
| 32 | Precuneus (Right) |

### 2.2 fMRI

All fMRI data were collected on a 3T Siemens Trio scanner with a 12-channel radio frequency coil. High-resolution T1-weighted structural images were acquired with a five-echo MPRAGE sequence with TE=1.64, 3.5, 5.36, 7.22, 9.08 ms, TR=2.53 s, TI=1.2 s, flip angle=7°, number of excitations=1, slice thickness=1 mm, field of view=256 mm, resolution=256×256. T2*-weighted functional images were acquired using a gradient-echo EPI sequence with TE=29 ms, TR=2 s, flip angle=75°, slice thickness=3.5 mm, slice gap=1.05 mm, field of view 240 mm, matrix size=64×64, voxel size=3.75 mm×3.75 mm×4.55 mm. An automated preprocessing pipeline and neuroinformatics system developed at the Mind Research Network (MRN, Scott et al., 2011) was used to preprocess the fMRI data. Five minutes of eyes-open resting data was collected from each participant.

After standard preprocessing (realignment, slice-timing correction, spatial normalization, and smoothing, see Houck et al., 2017), a subject-specific data reduction PCA was performed, retaining 100 principal components (PC). In order to use memory more efficiently, group data reduction was performed using an EM-based PCA algorithm and C=75 PCs were retained. The infomax algorithm (cf. Erhardt et al., 2011) was used for gICA from within the GIFT Toolbox (http://mialab.mrn.org/software/gift/). In order to estimate the reliability of the decomposition, the Infomax ICA algorithm was applied 10 times via ICASSO (Himberg et al., 2004) and the resulting components were clustered. Subject-specific maps and timecourses were estimated using a back-reconstruction approach based on PCA compression and projection (Calhoun et al., 2001, Erhardt et al., 2011). Of the 75 components returned by gICA, 39 were identified as BOLD-related based on their frequency content and spatial patterns (e.g. no edge-like effects, no components located in ventricles or white matter) [Allen et al., 2011].

To assess the frequency and structure of reoccurring FC patterns we applied the *k-*means clustering algorithm (Lloyd 1982) to windowed covariance matrices as in Allen et al. (2014), where dFNC was defined as the (Gaussian tapered) windowed zero-lag cross-correlations among reconstructed timecourses. We used a 22 TR window length (44 sec, following Miller et al., 2014 and Damaraju et al., 2014 and within the guidelines presented by Leonardi et al., 2015), slid 1 TR at each step, and computed pairwise correlations between time courses within these windows. Four “cluster states” were identified as optimal for k-means clustering using the silhouette and gap methods for the dFNC analysis. State transitions were computed for each subject at all windows. Temporal statistics of cluster states were calculated for each subject, including: frequency of occurrence (how often an individual visited a particular cluster state), dwell time (total time an individual remained in each cluster state), and number of transitions between cluster states. We summarized the temporal behavior of the resulting cluster states, which are then allowed to overlap in time, into meta-states; that is, a representation of how much a given subject is in each of the cluster states at each point in time. This approach builds distance vectors to the cluster centroids for each windowed FNC matrix. More specifically, windowed FNCs are modeled as “weighted sums of maximally independent connectivity patterns (CP)” (Miller et al., 2016). Discretized CP distance vectors are called meta-states. Global statistics were then calculated on the meta-states and compared between HC and SP groups, including: 1) The number of distinct meta-states subjects occupy during the scan length (“Number of states”); 2) The number of times that subjects switch from one meta-state to another (“Change between states”); 3) The range of meta-states subjects occupy, i.e., the largest L1 distance between occupied meta-states (“State span”); and 4) The overall distance traveled by each subject through the state space, i.e. the sum of the L1 distances between successive meta-states, (“Total distance”).

### 2.3 MEG

Five minutes of eyes-open resting state MEG data were acquired continuously. Artifact removal, correction for head movement, and downsampling to 250 Hz were conducted offline using Elekta Maxfilter software [Maxfilter, Elekta: (Taulu et al., 2004; Taulu and Simola, 2006)] with 123 basis vectors, a spatiotemporal buffer of 10 s, and a correlation limit of r=0.95. Cardiac and blink artifacts were removed using a signal-space projection (SSP) approach (Uusitalo and Ilmoniemi, 1997).

The cortical surface of each participant was reconstructed from T1-weighted MRI images using FreeSurfer for the automatic segmentation of the skull and scalp surfaces. Visual inspection confirmed that the automatic segmentation returned a reasonable solution. A repeatedly subdivided icosahedron was used as the spatial subsampling method which resulted in 10,242 locations per hemisphere. Coordinate system alignment was accomplished by first manually identifying fiducial landmarks and second by refining the alignment with the iterative closest-point algorithm (Besl and McKay, 1992) using the digitized scalp surface points. Source space analysis was conducted using dynamic statistical parametric mapping (MNE/dSPM), an anatomically constrained linear estimation approach (Dale et al., 2000). The regularization parameter was set to correspond to a signal-to-noise-ratio of 3 in the whitened data. Source orientation had a loose constraint of β=0.2, and a depth weighting of 0.8 was used. The forward solution was calculated using a single compartment boundary element method (BEM) (Hämäläinen and Sarvas, 1989; Mosher and Leahy, 1999); the surface was tessellated with 5120 triangles. Activity at each vertex of the cortical surface was mapped using a noise-normalized minimum norm estimate (Dale et al., 2000). In essence, MNE/dSPM identifies where the estimated current differs significantly from baseline noise (e.g., empty room data); this method also acts to reduce the location bias of the estimates (Gramfort et al., 2014). Spatial normalization was accomplished using FreeSurfer spherical coordinate system (Dale et al., 1999; Fischl et al., 1999) for group comparisons. Spatiotemporal source distribution maps downsampled to a 50 Hz sampling rate were obtained at each time point (providing an upper frequency bound of 25 Hz). Due to processing time and data storage considerations the first 60 seconds of each scan were projected into the brain volume and the files were converted to NIFTI format in order to be used with the GIFT toolbox.

Group spatial ICA (gsICA) was applied to the subject MNE/dSPM source-space maps using the GIFT toolbox (http://mialab.mrn.org/software/gift) as in our prior work (Houck et al., 2017). MEG ICA processing generally followed the procedures applied to the fMRI. Spatial maps were generated by decomposing the mixed MEG timecourses to yield a set of 32 spatially independent and temporally coherent networks. The final number of components was selected by determining that 1) networks (single or multiple areas of activation) were not being lost by the reduction in number of components and 2) that the same area was not being “broken up” into numerous components when only a single area of activation was present (See Fig. 1 for examples of components). This was an important consideration since later analyses involved multiple statistical comparisons. Furthermore, unlike with fMRI, this data should contain a minimum of artifact components (in the present case we found no artifact components) due to noise reduction and artifact removal during preprocessing. As with fMRI, subject-specific maps and timecourses were estimated using a back-reconstruction approach based on PCA compression and projection (Calhoun et al., 2001; Erhardt 2011; Houck et al., 2017). Although a beamformer source-space analysis technique has been used in a similar manner before (as input to an ICA, Houck et al., 2017), we show here that the MNE/dSPM measure used in this way gives reasonable, and arguably more focal (although not point-like) components. This may be due to the differential sensitivity of MNE-based methods to connectivity in non-ERP data (Hincapié et al., 2017). Furthermore, simulation has shown that MNE provides better connectivity estimation than beamformers if the interacting sources are simulated as extended cortical patches with high within source coherence (Hincapié et al., 2017). Which method to use is also an important consideration when correlations between sources may lead to partial or full signal cancellation in the beamformer. Lastly, a k-means clustering, dFNC, and meta-state analysis was conducted between all 32 components (Table 1), following the same procedure described above for fMRI.

**Figure 1.**
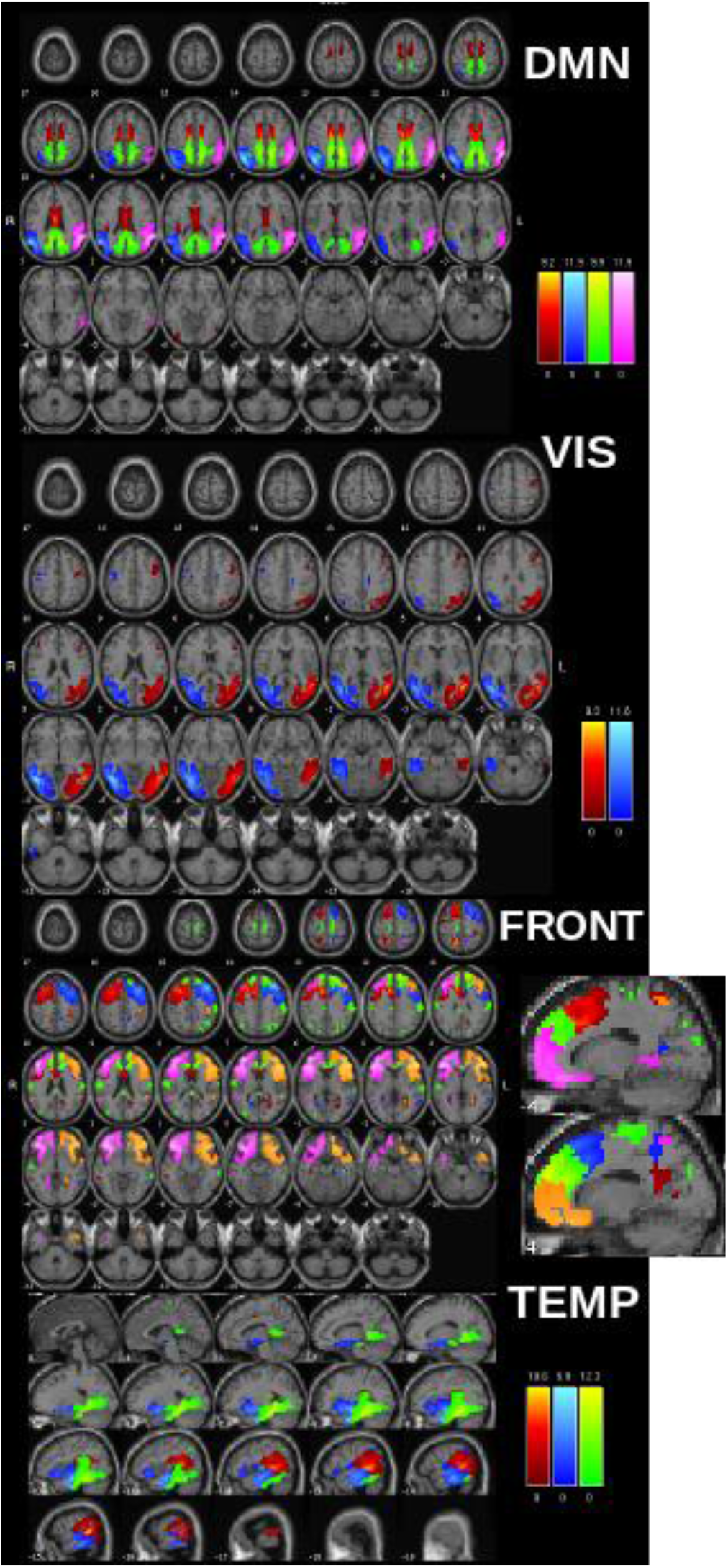
ICA component maps from resting MEG data separated into anatomic domains. Note that each color within each network represents a different independent component (IC). Please note that the same color in a different network does not necessarily represent the same region. DMN: Pink & Blue=inferior parietal/angular gyrus; Green=cingulate (isthmus); Red=posterior cingulate; VIS: visual components. FRONT: Primarily frontal components (some overlap with cingulate areas). TEMP: Primarily temporal components.

One question which needed to be addressed was the length of the window to use for the dFNC analysis. For fMRI we chose to follow a well-established precedent of a 22-TR window (= 44 sec) (Allen et al., 2014). Such a long time window was not considered for the MEG data since we were interested in investigating the faster oscillations available from this modality. We therefore chose a 4 sec window length for the analysis. This allowed us to capture our range of frequencies of interest (1 Hz – 25 Hz) giving us the ability to see connections at higher frequencies unavailable via the fMRI analysis while avoiding “washing out” information from higher frequencies with too long a window.

In addition, the decision of how many cluster states to use in the MEG dFNC analysis presented some questions. First was whether the MEG dFNC should closely parallel the fMRI dFNC steps. This was abandoned for two reasons: 1) due to the temporal resolution of the MEG data there was no a priori reason to expect that the two modalities would reveal the same number of states, and 2) different quantities of data were evaluated, with 60 seconds of 50 Hz data in the MEG case (i.e., 3000 sampling points), as opposed to 300 seconds with a 2-sec TR in the fMRI (i.e., 150 sampling points). We initially used the gap and silhouette methods to estimate the number of k-means cluster states, however we found that due to a single outlier individual the 2 cluster state suggestion was unstable. Investigation of 3, 4, and 5 cluster states indicated that 4 states was optimal for the present dataset, as this gave stable, replicable results which consistently grouped all the data from the single outlier individual into its own cluster state, and also gave reasonable occupancy percentages for the remaining 3 states (i.e. the percentage of correlation maps in each meaningful state were: 19%, 64%, 17%). The state which contained the outlier did not return any statistics for the group analysis since it did not contain sufficient data (i.e. only a single individual entered that state and all data for that individual was associated with that state), this state was removed from all additional analyses and did not influence any of the results shown.

Lastly, we summarized the temporal behavior of the resulting cluster states, which are now allowed to overlap in time, into meta-states; that is, a representation of how much a given subject is in each of the cluster states at each point in time. Global statistics were then calculated on the meta-states and compared between HC and SP groups. Recall these are: “Number of states”, “Change between states”, “State span”, and “Total distance”.

## 3. Results

### 3.1 dFNC and Meta-State statistics of fMRI data

We found that 4 k-means cluster states characterized the temporal dynamics of a 5 min eyes-open resting state fMRI scan, for both HC and SP, with a minimum of 24 participants entering each state at some point during the 5 mins of data collection (Fig 2.A). Furthermore, we found that for some cluster states SP and HC spent significantly different amounts of time there, as determined by dwell time. Particularly, SP visited and remained in state 1 for a significantly longer time than HC, whereas for state 3 the trend was reversed, with HC remaining in that state for a significantly longer time (**Fig 2.B**). On average, schizophrenia patients spent less time than healthy controls in a state typified by strong, large-scale connectivity (state 3, many correlation showing r > 0.5). No significant inter-individual variability between groups was found for any cluster state.

**Figure 2.**
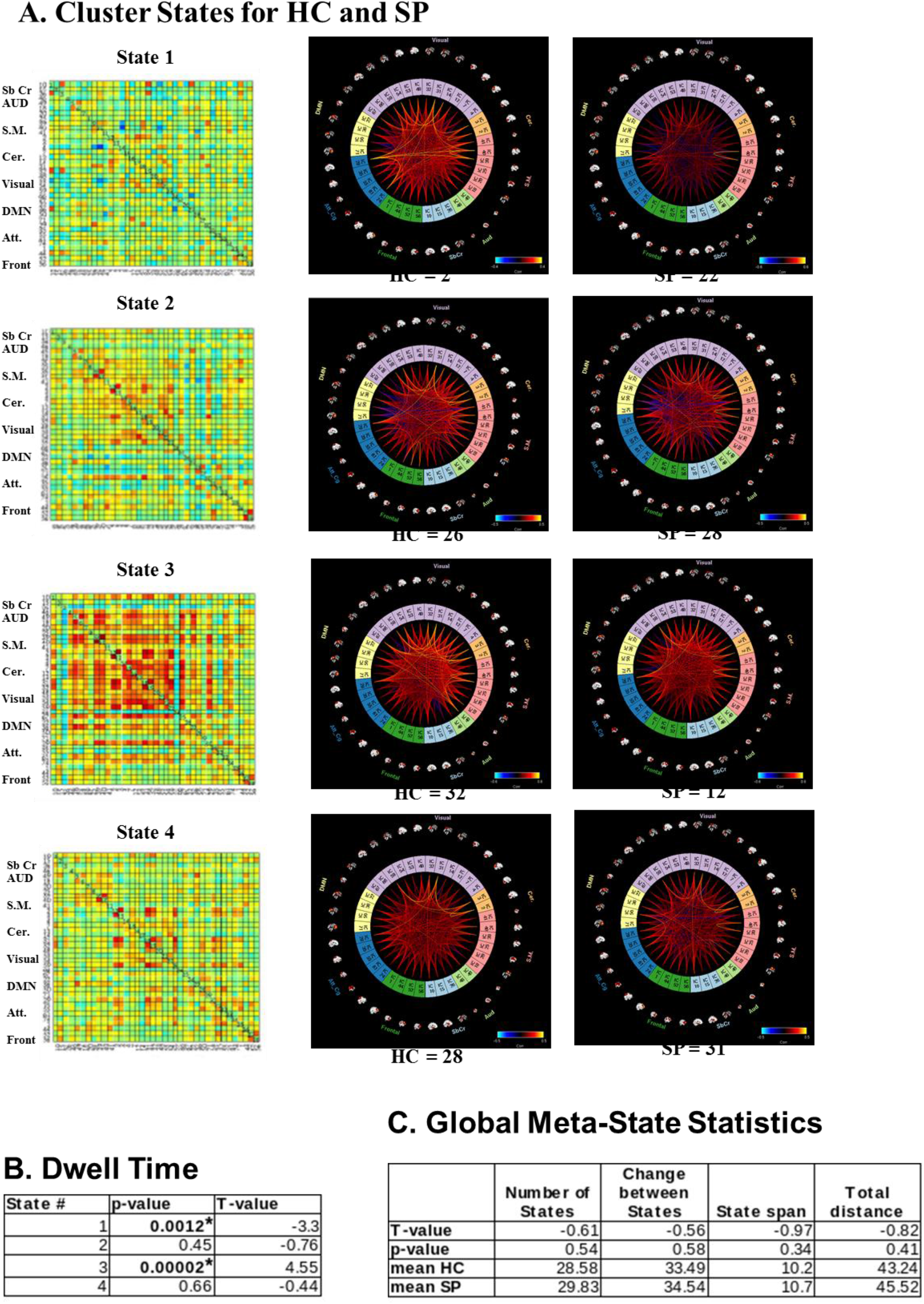
A. fMRI k-means cluster states derived from ICA components, for HC and SP groups, number of participants that entered each state is indicated at the bottom of each plot. Color-coded as follows: light blue=subcortical, light green=auditory, pink=sensory motor, orange=cerebellar, light purple=visual, yellow=DMN, blue=attention, dark green=frontal. Insets show view of dFNC as correlation grids (components x components) in same network order (top to bottom). B. Average amount of time HC and SP spend in each state, i.e. dwell time. C. Global meta-statistics for HC vs SP groups. All t-tests represent HC-SP. ‘*’ indicates significance at p < 0.05, FDR corrected.

We also investigated global statistics, and for these meta-state statistics, found no significant group differences for any measures including: number of states, change between states, state span, nor total distance (as defined in “Methods”).

### 3.2 dFNC and Meta-State statistics of MEG data

We found that 3 k-means cluster states were stable and characterized the temporal dynamics for 1 min of eyes-open resting MEG data, for both HC and SP, where a minimum of 58 participants entered each state at some point during the 1 min of data investigated (**Fig 3.A**). We found a significant overall between-group difference for state 2, with more variability for SP than for HC (i.e. a significant difference was observed in inter-individual variability for how closely each group resembled the state).

**Figure 3.**
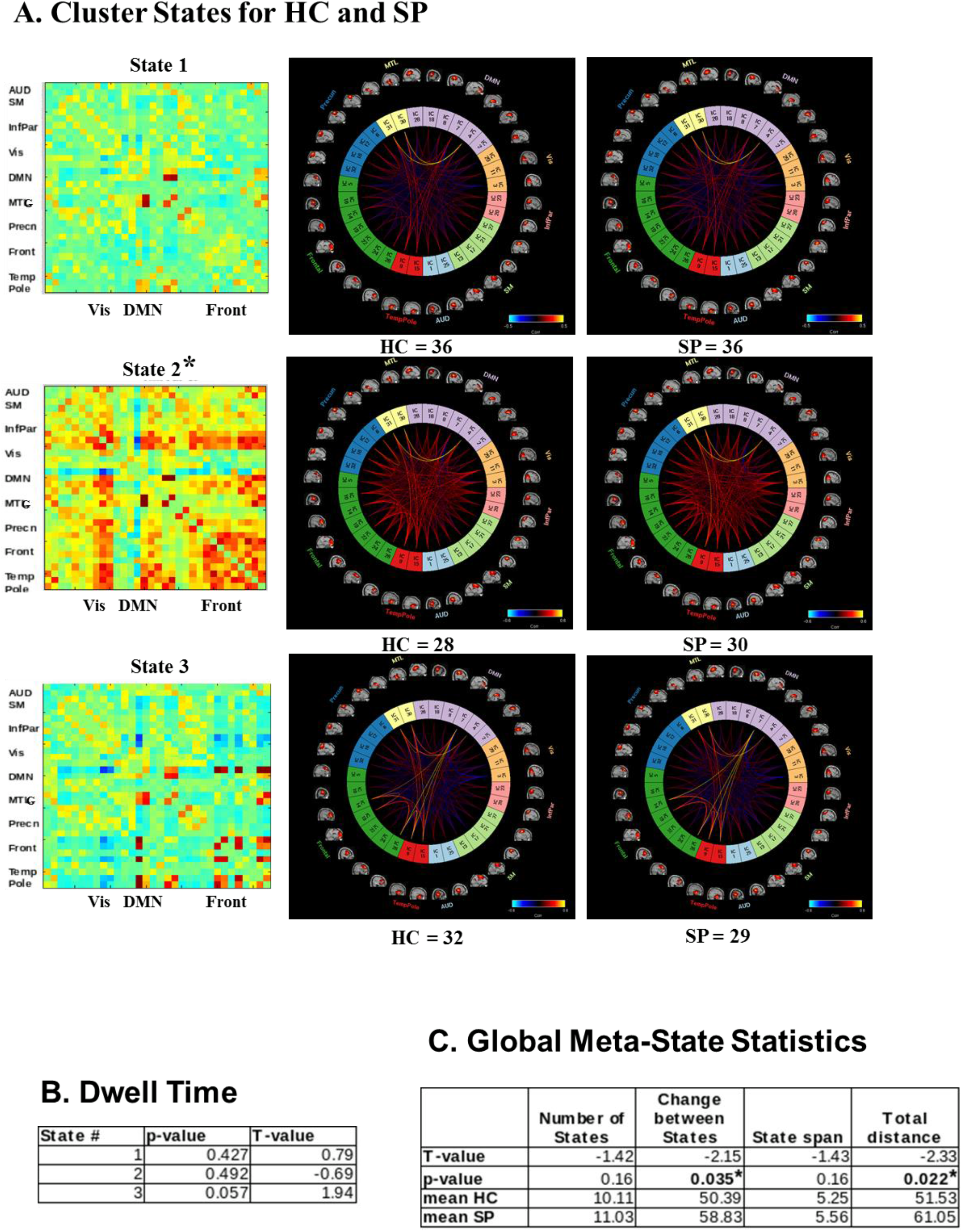
A. MEG k-means cluster states derived from ICA components, for HC and SP groups, number of participants that entered each state is indicated at the bottom of each plot. Color-coded as follows: light blue=auditory, light green=sensory motor, pink=inferior parietal, orange=visual, light purple=DMN, yellow=MTL, blue=precuneus, dark green=frontal, red=temporal pole. Insets show view of dFNC as correlation grids (components x components) in same network order (top to bottom). B. Average amount of time HC and SP spent in each state, i.e. dwell time. C. Global meta-statistics for HC vs SP groups. All t-tests represent HC-SP. ‘*’ indicates significance at p < 0.05, FDR corrected.

In contrast to the fMRI results we found no group differences in dwell time between HC and SP groups (**Fig 3.B**). However, in this data we see group differences in the global meta-state statistics “change between states” and “total distance” (**Fig 3.C**).

## 4. Discussion

### 4.1 dFNC of fMRI and MEG: Do they provide complementary information?

Our fMRI results, which reveal that SP spent significantly less time in a highly connected cognitive state (Dwell time, state 3 **Fig. 2B**), parallel and replicate those of Damaraju et al. (2014) although we investigated an eyes-open resting state in contrast to the eyes-closed paradigm used in their study. However, for meta-state statistics calculated from fMRI data we found no significant group differences for any measures including: number of states, change between states, state span, nor total distance (as defined in “Methods”). This is in contrast to Miller et al. (2014 and 2016) who introduced and investigated these statistics between HC and SP in eyes-closed resting data and found significant “reduced connectivity dynamism” in all statistics for SP relative to HC for these global statistics. Of additional note is that the directionality (i.e. which group, HC or SP, showed the higher mean value) of all four global meta-state statistics is in the opposite direction as was found in Miller et al. (2014), with HC exhibiting less movement between states than SP. However, we again point out that an eyes-open resting state was used for the current study and with a smaller sample size, whereas Miller used a eyes-closed resting state, both potentially effecting which states were detected and how frequently they are visited (McAvoy et al., 2012).

Our MEG dFNC results revealed a significant overall between-group difference for state 2, with more variability for SP than for HC (i.e. significant difference was observed in inter-individual variability for how closely each group resembled the centroid of the cluster state). Because this state includes many high correlations between fronto-frontal and fronto-parietal regions (i.e. *r > 0.5*), this group difference is in keeping with numerous results that indicate that there is a frontal dysfunctional connectivity (i.e., dysconnectivity; Bullmore, Frangou, & Murray, 1997; Friston & Frith, 1995) for SP, a dysconnectivity seen both among frontal regions, e.g. insula and lateral frontal cortex (Palaniyappan et al., 2013) and between frontal regions and more distal areas, particularly fronto-parietal control network regions (Wu et al., 2017; Roiser, et al., 2013). We also saw a strong correlation between activity in bilateral medial temporal gyrus (MTG) and parahippocampus for all cluster states and groups, a relationship that may deserve additional investigation due to their involvement in memory, particularly recollection (Eichenbaum et al., 2007), as well as to everyday functioning in SP (Hanlon et al., 2012). It appears that the timescale of the MEG data may be attuned for such study.

In contrast to the fMRI results we found no group differences in dwell time between HC and SP groups (**Fig 3.B**). However, it is in this data where we see group differences in some of the global meta-state statistics, specifically the “change between states” and the “total distance” (**Fig 3.C**). Recall these are the number of times that subjects switch from one meta-state to another and the overall distance traveled by each subject through the state space (the sum of the L1 distances between successive meta-states), respectively. This indicates that not only are SP changing states more often than HC, but they are also changing to states that are *more different in comparison to the previous state occupied*. Interestingly, and something which should be investigated further, we find that in all cases global meta-state statistics occur in the same direction even when failing to reach significance. In other words, SP show higher mean values for meta-state statistics relative to HC for both the MEG and fMRI analyses in the current study, which is reversed from what was found in Miller et al. (2014) for an eyes-closed resting state.

In the present study we found that the information that can be gained from a dFNC analysis of resting state data of the same participants differs between the neuroimaging modalities of fMRI (differences seen in cluster state level statistics) and MEG (differences seen mostly in global-level meta-state statistics). Therefore the fusion of these modalities is expected to reveal additional information (Calhoun and Liu, 2016), potentially informing identification/classification (e.g. individuals may have co-morbidities), new treatment targets, and/or potential treatment success. The next step will include determining the best way of combining these data within an overarching framework to maximize the information from each modality and their combination (e.g. joint-ICA). The major limitations of the current study was the use of only 1 min of MEG data for the analyses, and the downsampling of the MEG data to 20ms necessitated by computing time and memory constraints, which resulted in an upper frequency limit of 25Hz, and therefore an inability to probe gamma band relationships at this time. We are currently working on implementing a faster method for the analysis and will soon have hardware supplying additional memory resources alleviating these shortcomings. Arguably, one could travel to a greater number of distinct “states” were one to have more time to do so, although of course both HC and SP had the same limitation. In addition, were this the case we would argue that many more than 3 viable states would have been detected in the 1 min of MEG data analyzed. A higher sampling rate could plausibly reveal more interactions and transitions (including gamma band), however a preliminary analysis with a 10ms sampling rate (50Hz maximum frequency) reveled similar results to those presented here. It has been conjectured that even 5 mins of data (an often used scan duration for fMRI) may be too short for dFNC determined from fMRI data. Indeed, improvements in the reliability of resting state data tend to rise with scan length, plateauing at a scan length of 13 minutes (Birn et al., 2013). Consistent with this finding, Liuzzi, et al. (2016) showed that large improvements in repeatability were apparent when using a 10 min, compared to a 5 min recording for MEG resting state functional connectivity analyses. However, that study was looking for a single “canonical” state, whereas the present study, using windowed correlations and k-means clustering is evaluating a set of states whose number is data driven, which may have a different, perhaps shorter, optimal scan duration. It has been shown that FNC states in electrophysiological data can be quite transient (100–200 ms), suggesting that the resting brain is changing between different patterns of repeated activity at a rapid pace at the neuronal temporal scale (Vidaurre et al., 2016). Clearly the time scales of these states need further investigation. However, regardless of the time scale, imaging modality, or underlying biological signal, the primary finding across studies, including the present study, is that d/FNC evolves as a multi-stable process passing through multiple and reoccurring discrete cognitive states, rather than varying in a more continuous sense (Cabral, et al., 2016; Hutchison et al., 2013; Allen et al., 2014; Hansen et al., 2015; Preti et al., 2016).

### 4.2 Conclusions

Our results here show how two neuroimaging modalities, MEG and fMRI, can each reveal distinct differences between HC and SP groups, and how the differences emerge in different metrics for each modality. For the eyes-open resting state investigated we found group differences at the “cluster-level” for fMRI (i.e., dwell time), whereas for MEG we found differences at what has been referred to as the “global-level” (i.e., change between states and distance traveled: Miller et al., 2014). This indicates the importance of future work which usefully combines these two neuroimaging methodologies, taking advantage of the distinct information contained in each, in order to e.g. better differentiate clinical populations with overlapping symptoms, for example using a joint-ICA. Future work will also aim to identify which metrics correlate with illness characteristics such as symptoms, chronicity, and cognitive impairment. In addition we presented a novel MEG analysis pipeline, which incorporates a source-space analysis (MNE/dSPM) as input to a group ICA, the networks/components of which may then be used for a dFNC analysis.

## Acknowledgments

Research reported in this publication was supported by the National Institute on General Medical Sciences, National Institute on Alcohol Abuse and Alcoholism, and National Institute of Biomedical Imaging And Bioengineering of the National Institutes of Health under award numbers P20GM103472, K01AA021431, R01EB006841, and R01REB020407. Additional support was provided by NSF grant 1539067. The content is solely the responsibility of the authors and does not necessarily represent the official views of the National Institutes of Health or the National Science Foundation.

